# Hyperactivated YAP1 Drives an Invasive EMT Subtype of Cervical Squamous Cell Carcinoma

**DOI:** 10.64898/2026.01.18.700207

**Authors:** Jiyuan Liu, Xiangmin Lv, Jinpeng Ruan, Chunbo He, Guohua Hua, Peichao Chen, Xingeng Zhao, Davie Shi, Siyi Yang, Anneta Polakkottil, Madelyn L Moness, Brianna Carbajal, Gunjan Kurma, Reema Guda, Ayan A Mohamed, Olukemisola Abioye, Ruslan Sadreyev, John S. Davis, Cheng Wang

**Author notes:** **Corresponding Author:** Cheng Wang, Ph.D., Vincent Center for Reproductive Biology Department of Obstetrics and Gynecology Massachusetts General Hospital, Harvard Medical School Boston, MA 02114. Co-first authors (Jiyuan Liu, Xiangmin Lv).

## Abstract

Cervical cancer (CVC) is classically understood as an HPV-driven disease in which high-risk HPV infects cervical basal epithelial cells, induces neoplastic transformation, and drives upward epithelial expansion, pathogenic events underlying the success of cytology- and HPV-based screening. With widespread HPV vaccination and screening, the WHO has launched the CVC elimination initiative and believes that CVC will be the first cancer that would be eliminated as a public health problem. However, statistics showed that CVC remains the most common gynecologic malignancy worldwide, and in the United States, where HPV testing and Pap smears have reached near-maximal implementation, the CVC mortality rate has plateaued for more than two decades. These epidemiologic trends suggest the existence of a subset of CVC that can escape current screening strategies. Here, we demonstrate that hyperactivation of YAP1 caused by disruption of Hippo-YAP signaling is sufficient to induce a subtype of HPV-independent invasive CVC that lacks surface lesions and therefore evades HPV- and cytology-based detection. Using single-cell RNA sequencing combined with high-resolution spatial transcriptomics, we define the cellular architecture, molecular pathways, and immune microenvironment underlying this invasive subtype. We show that YAP1-driven tumors adopt an EMT-high transcriptional state and selectively recruit immunosuppressive myeloid-derived suppressor cells that functionally interact with cancer cells to promote invasion and progression. Our findings suggest that targeting the disrupted Hippo signaling may offer a new strategy to prevent the subset of invasive CVC that current HPV- and Pap-based programs cannot detect, an essential step toward achieving global CVC elimination.

## INTRODUCTION

Cervical cancer (CVC) arises from the epithelial lining of the uterine cervix. It is widely accepted that development of CVC is tightly linked to the infection of high-risk human papillomavirus (hrHPV). Currently, hrHPV is recognized as a necessary causal agent for virtually all cases of CVC (1, 2). Molecular studies have shown that hrHPV infects the basal cells of the cervical transformation zone, leading to deregulated expression of the viral oncogenes E6 and E7, which drive the neoplastic progression from low-grade squamous intraepithelial lesions (LSIL) to high-grade lesions (HSIL) and, eventually, to carcinoma in situ and invasive cancer (2).

The introduction of the Pap test revolutionized CVC prevention by enabling detection of precancerous cytologic abnormalities, dramatically reducing incidence and mortality where screening programs became widespread (3, 4). Building on the HPV-driven model of cervical carcinogenesis, modern prevention strategies now include HPV DNA-based screening and prophylactic HPV vaccination, the latter of which has shown high efficacy in preventing persistent HPV 16/18 infection and cervical intraepithelial neoplasia (CIN) in clinical trials (5–7). These advances led to international consensus that CVC could become the first human cancer to be eliminated at the population level, culminating in the launch of the World Health Organization’s global elimination initiative (8–13).

Despite these public health achievements, recent epidemiological data underscore that CVC remains a significant cause of cancer morbidity and mortality worldwide. According to the latest GLOBOCAN 2022 analysis, approximately 662,000 women developed cervical cancer and 349,000 died of the disease in 2022, confirming that CVC continues to rank among the most common cancers in women globally (14). Even in high-resource settings such as the United States, where Pap testing and HPV screening have long been widely implemented, progress in reducing mortality has stalled over the past two decades with recent analyses showing minimal improvement in national mortality trends (15). These population-level observations point to the possibility that current prevention strategies, while highly effective against HPV-driven precancers, may not detect all biologically relevant pathways to cervical carcinogenesis. Such evidence also raises an important biological and clinical question: are additional, HPV-independent mechanisms contributing to cervical cancer initiation and early progression?

Interestingly, our recent work demonstrated that disruption of Hippo signaling is sufficient to drive cervical cancer in the absence of HPV, establishing a mechanistic basis for HPV-independent CVC initiation (16). Here, we showed that hyperactivation of YAP1 in cervical epithelial cells induces early invasive cancer cell populations capable of breaching the stromal boundary during the initial stages of carcinogenesis. Crucially, the biology of these early invasive lesions, rapid invasion, lack of HPV involvement, and limited cytologic atypia, renders them undetectable by conventional Pap cytology and by HPV-based molecular screening. These findings identify a previously unrecognized subtype of cervical cancer and provide a biological explanation for persistent mortality despite comprehensive HPV-focused screening programs.

## RESULTS

### Hyperactivated YAP1 is closely associated with CVC invasiveness and poor patient outcome

Thanks to the effectiveness of HPV- and Pap test-based screening, many cases of CVC could be diagnosed at an early stage, resulting in a fairly good prognosis. However, a subset of CVC is featured with late stage at presentation (even after regular pap tests), distant metastasis, and very poor prognosis, suggesting that a subset of CVC can successfully evade from current screening system. To investigate the existence of this subset and cellular and molecular mechanisms contributing to their late presentation and poor prognosis, we analyzed the clinical, genomic, transcriptomic, and proteomic datasets generated by the Cancer Genome Atlas (17). We found that once patients surpass the first ∼22 months following diagnosis, approximately 70% remain disease-free for more than 200 months (Fig. 1A). Based on this finding, we divide patients into two groups: patients deceased within 22 months of diagnosis (“<22-mo”) and patients survived beyond 22 months (“>22-mo”). The overall survival (OS), progression-free survival (PFS), disease-specific survival (DSS), and disease-free survival (DFS) in >22-mo groups were significantly higher than those of patients in <22-mo group. The DEG analyses showed that both YAP1 mRNA and protein levels were significantly higher in the <22-mo group compared with the >22-mo group (Fig. 1B). Consistent with these findings, the two groups exhibited markedly different YAP1 copy number variation (CNV) distributions (Pearson’s Chi-squared test, *p* = 4.2 × 10⁻⁹) (Fig. 1C). High-level YAP1 amplification occurred in roughly half of the patients who died within 22 months, but in only ∼10% of long-term survivors. Gene Set Enrichment Analysis further demonstrated that YAP1 transcriptional activity was strongly enriched in the <22-mo group (Fig. 1D), and proteomic profiling revealed significantly elevated YAP1 signature scores in this same group (Fig. 1E). Together, these findings support YAP1 activation as a molecular feature of poor prognosis in CVC.

**Figure 1.**
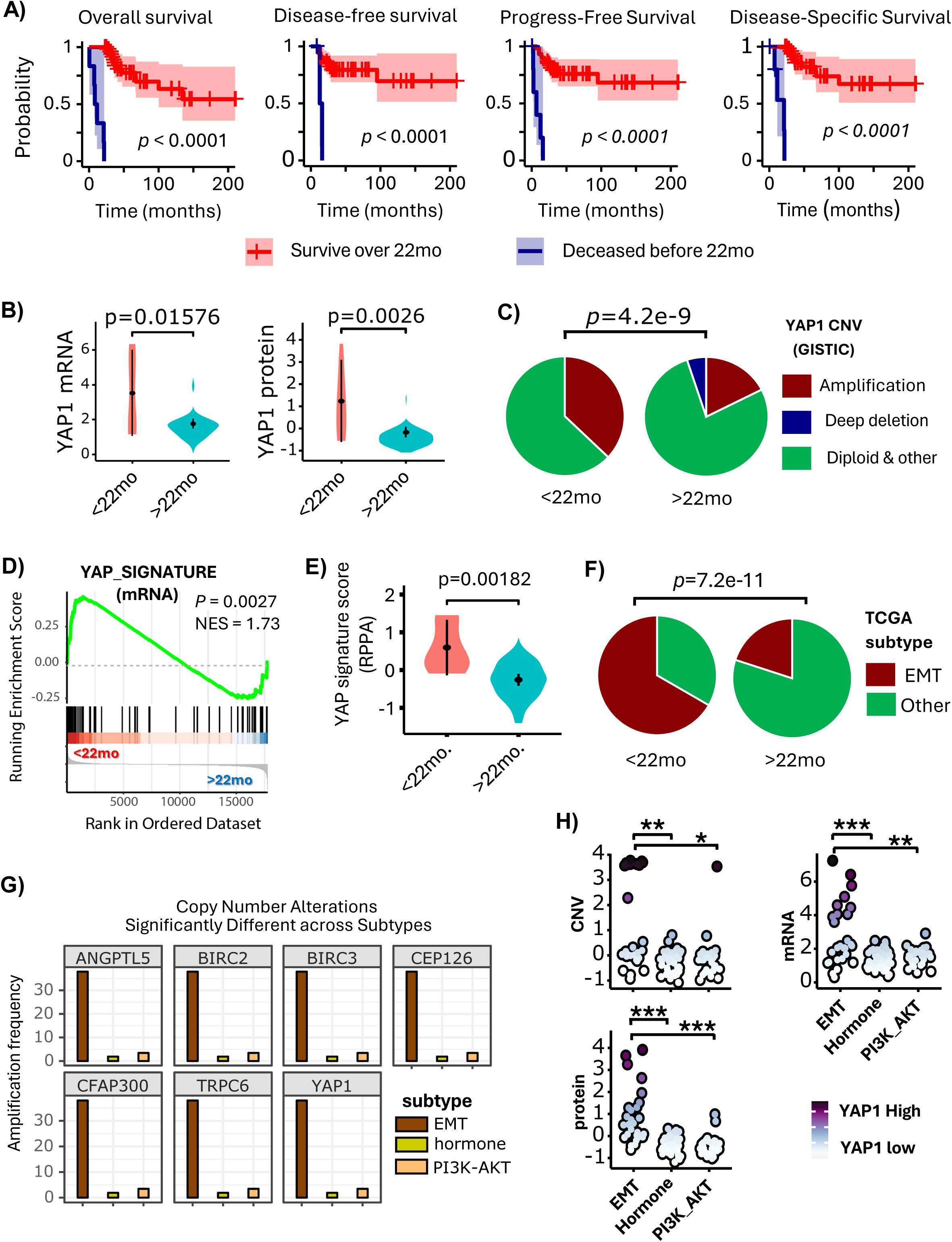
YAP1 is hyperactivated in a portion of cervical cancer patients that have poor prognosis. **A**) Overall survival (OS), Progress-free survival (PFS), Disease-specific survival (DSS) and Disease-Free survival (DFS) of cervical cancer patients who survived over 22 months (n=144) and those who deceased within 22 months (n=40). *P* values were derived from log rank test. **B**) Expression of *YAP1* mRNA and protein in cervical cancer patients who deceased within 22 months (<22 mo) and those who survived over 22 months (>22 mo). *P* values were from unpaired two-sided t test. **C**) YAP1 copy number variation (CNV) in cervical cancer patients who deceased within 22 months (<22 mo) and those who survived over 22 months (>22 mo). Data is presented as GISTIC values. *P* values were from Pearson’s Chi-squared test. **D**) Gene Set Enrichment Analysis (GSEA) plot showing enrichment of YAP1 signature genes in patients who deceased within 22 months (<22 mo) when compared to those who survived over 22 months (>22 mo). *P* values were derived from GSEA permutation based on an adaptive multi-level split Monte-Carlo scheme implemented in fgsea R package. **E**) Relative protein levels (RPPA score) of YAP1 signature genes in cervical cancer patients who deceased within 22 months (<22 mo) and those who survived over 22 months (>22 mo). p values were from unpaired two-sided t test. **F**) Frequency of cervical cancer subtypes (classified using TCGA cervical cancer subtyping strategies) in cervical cancer patients who deceased within 22 months (<22 mo) and those who survived over 22 months (>22 mo). *P* values were from Pearson’s Chi-squared test. **G**) A comparison of amplification frequency of *YAP1* and its downstream genes (*ANGPTL5*, *BIRC2*, *BIRC3*, *CEP126*, *CFAP300*, and *TRPC6*) across EMT, Hormones, and PI3K-AKT subtypes. q values are Benjamini-Hochberg adjusted p values derived from Chi-squared test. A *q* values < 0.05 was considered as significantly different when compared to each other. **H**) A comparison of copy number variation (CNV), mRNA expression, and protein levels of YAP1 across EMT, Hormones, and PI3K-AKT subtypes. *p* values were from unpaired two-sided t test. A *p* values < 0.05 was considered as significantly different. * *p*<0.05, ** *p*<1e-3, *** *p*<1e-5, when compared to each other.

Based on deep sequencing data, TCGA classified CVC into three subtypes, namely, Hormone, PIK3-AKT, and EMT subtypes, in which EMT subtype was featured with high expression of YAP1 and poor prognosis (17, 18). Consistent with TCGA findings, we found that the majority of cases in the <22-mo group could be categorized as EMT subtype based on TCGA gene signature (Fig. 1F). Cross-subtype genetic analyses also demonstrate that genes encoding for *YAP1* and the major tumor-promoting components of the Hippo-YAP signaling pathway, including, but not limited to *ANGPTL5*, *BIRC2/3*, *CEP126*, *CFAP300*, and *TRPC6*, are all amplified and highly expressed in the EMT subtypes (Fig. 1G & Fig. 1H). Strikingly, we also found that CVC samples with YAP1 CNV > 2, log2 YAP1 mRNA > 3, or RPPA-derived YAP1 Z-score > 1.5 would be exclusively classified as EMT subtype (Fig. 1H). Since EMT is tightly associated with tumor invasiveness (19–21), these observations imply that the hyperactivation of YAP1 is closely associated with CVC invasiveness and poor survival outcome.

### Hyperactivation of YAP1 in cervical epithelial cells drives development of HPV-independent invasive cancer in vivo

To test the hypothesis that hyperactivated YAP1 in the cervical epithelium contributes to CVC invasiveness, we generated a transgenic mouse model expressing a constitutively active YAP1 (YAP^S127A^) in cervical epithelial cells under the control of the *Krt14* promoter and induction of doxycycline (Dox) (“*Krt14*-*rtTA*;*Tet*-on-*YAP*^S127A^”, hereafter refereed as KY mice). Estradiol-17β pellets (0.25 mg, 90-day release) were implanted as reported previously to promote tumorigenesis (16). Dox (0.03 mg/mL in drinking water) was used to induce transgene expression(16). KY mice received water without Dox were used as negative control. Wild-type mice and krt14-rtTA mice treated with the same dose of Dox were used as additional negative controls. Early stage tumors developed in the transformation zone of uterine cervix within 2 months of Dox induction (Fig. 2A, 2B) with a portion of KY mice developed ureteral obstruction, a feature that is also observed in human patients with advanced stage invasive cervical cancer (22). No obvious phenotype was observed in the control groups. Histological analyses showed that cancer induced by hyperactivated YAP1 is extremely invasive. The newly formed YAP1-high tumor cells are so invasive that they directly break through the basal membrane and invade into the stromal layer, leaving the lumen epithelia intact (Fig. 2B, 2C, and 2D). The relatively low expression of E-cadherin in the foci of invasive cancer in the stromal layer, which was detected by both immunofluorescence (Fig. 2G), suggest that these are cervical carcinoma cells, not cells from the stroma. Since these tumors are HPV-independent and they invade inward stroma since cancer initiation, this subtype of invasive cancer cannot be screened by current pap test- and HPV-based screening system.

**Figure 2.**
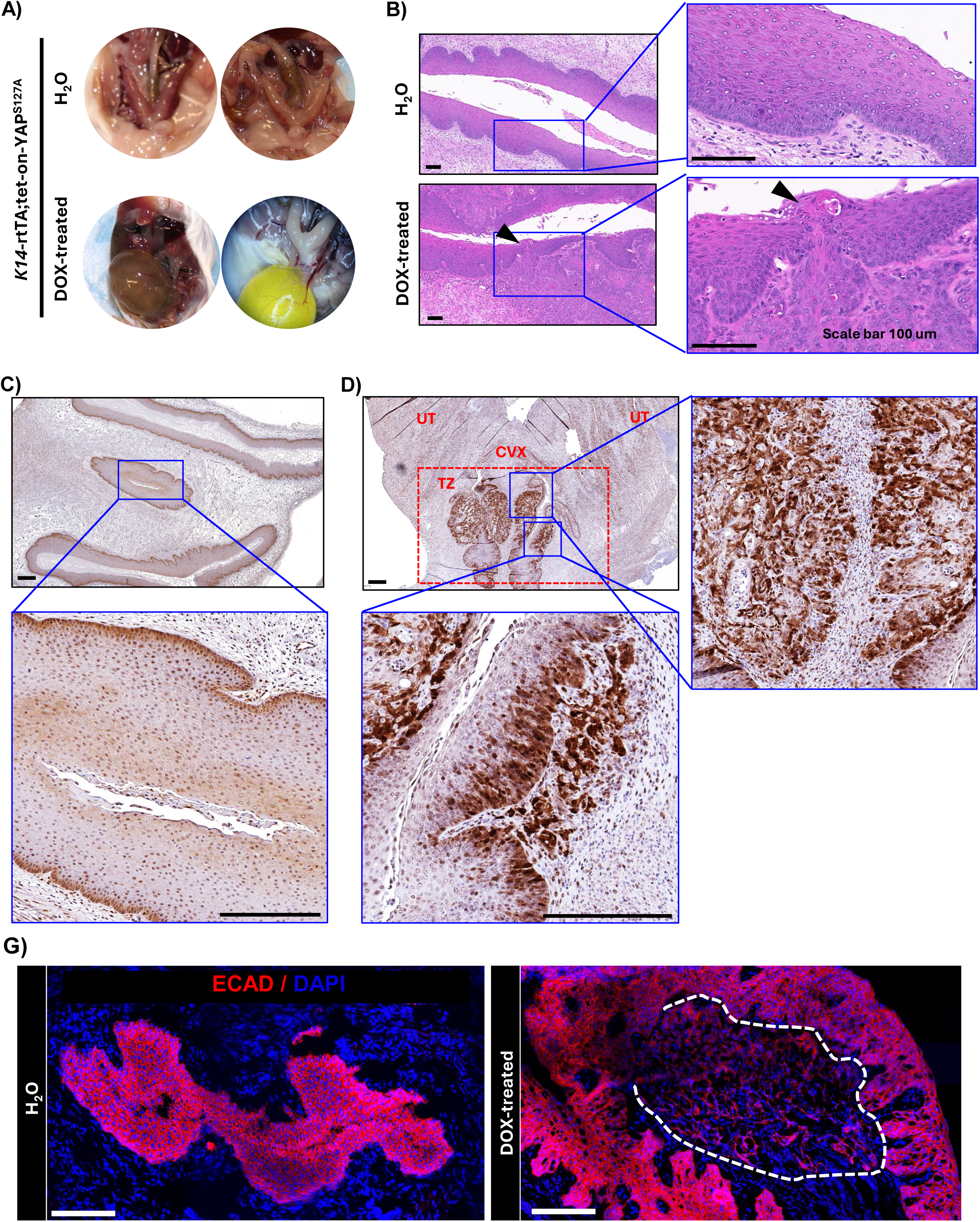
Activation of YAP1 in cervical epithelial cells leads to tumors exhibiting high invasiveness and strong EMT signatures. **A**) Representative images showing the ureter obstruction in of *Krt14*-rtTA;Tet-on-YAP^S127A^ mice induced by doxycycline (DOX-treated, lower panel). *Krt14*-rtTA;Tet-on-YAP^S127A^ mice administered with water (H_2_O, upper panel) were used as control. Note the enlarged bladders caused by the ureteral obstruction due to tumorigenesis in DOX-treated *Krt14*-rtTA;Tet-on-YAP^S127A^ mice induced (n = 5-8/group). **B**) Representative images showing the histology (H&E staining) of cervical epithelium in the *Krt14*-rtTA;Tet-on-YAP^S127A^ mice induced with doxycycline (DOX, lower panel) or water (H2O, upper panel). Arrowheads point to the invading site where transformed epithelial cells break the basal membrane and invade into the stromal layer of cervical tissues. Scale bar: 100 µm. **C**) representative images showing the expression of YAP1 protein (in brown, detected by IHC) in cervical tissues of *Krt14*-rtTA;Tet-on-YAP^S127A^ mice induced with water as control. Scale bar: 200 µm. **D**) representative images showing the expression of YAP1 protein (in brown, detected by IHC) in cervical tissues of *Krt14*-rtTA;Tet-on-YAP^S127A^ mice administered with doxycycline. Scale bar: or DOX (lower panel). A dash-lined red rectangle box was used to highlight the transformation zone (TZ) of mouse uterine cervix. Two blue-line squares in the transformation zone highlight the newly formed cancer tissues in the cervix of Dox-induced *Krt14*-rtTA;Tet-on-YAP^S127A^ mice. UT: uterine tubes; CVX: cervix. Scale bar: 200 µm. **E**) Representative images showing expression of E-cadherin (in red, detected by fluorescent immunohistochemistry) in the cervical tissue of *Krt14*-rtTA;Tet-on-YAP^S127A^ mice administered with water (H2O, left) or doxycycline (Dox-treated, right). Carcinoma cells invaded into the stromal area, which express relatively low E-cadherin (CDH1^+^), were circled with a white dished-line. Scale bar: 100 µm.

### Single-cell RNA sequencing reveals cellular landscape alterations during development of YAP1-driven invasive cervical cancer

To characterize cellular and molecular changes accompanying YAP1-driven cervical carcinogenesis, we performed single-cell RNA sequencing (scRNA-seq) on cervical tissues from 17β-estradiol–implanted KY mice treated with or without doxycycline following a protocol developed in our lab (23). Unsupervised clustering using the Seurat v5 workflow (24) together with established cervical epithelial and stromal marker genes (25, 26) identified nine major cell populations, including basal/parabasal epithelial cells, keratinocytes, cancer cells, endothelial cells, lymphocytes, myeloid cells and stromal cells (Fig. 3A).

**Figure 3.**
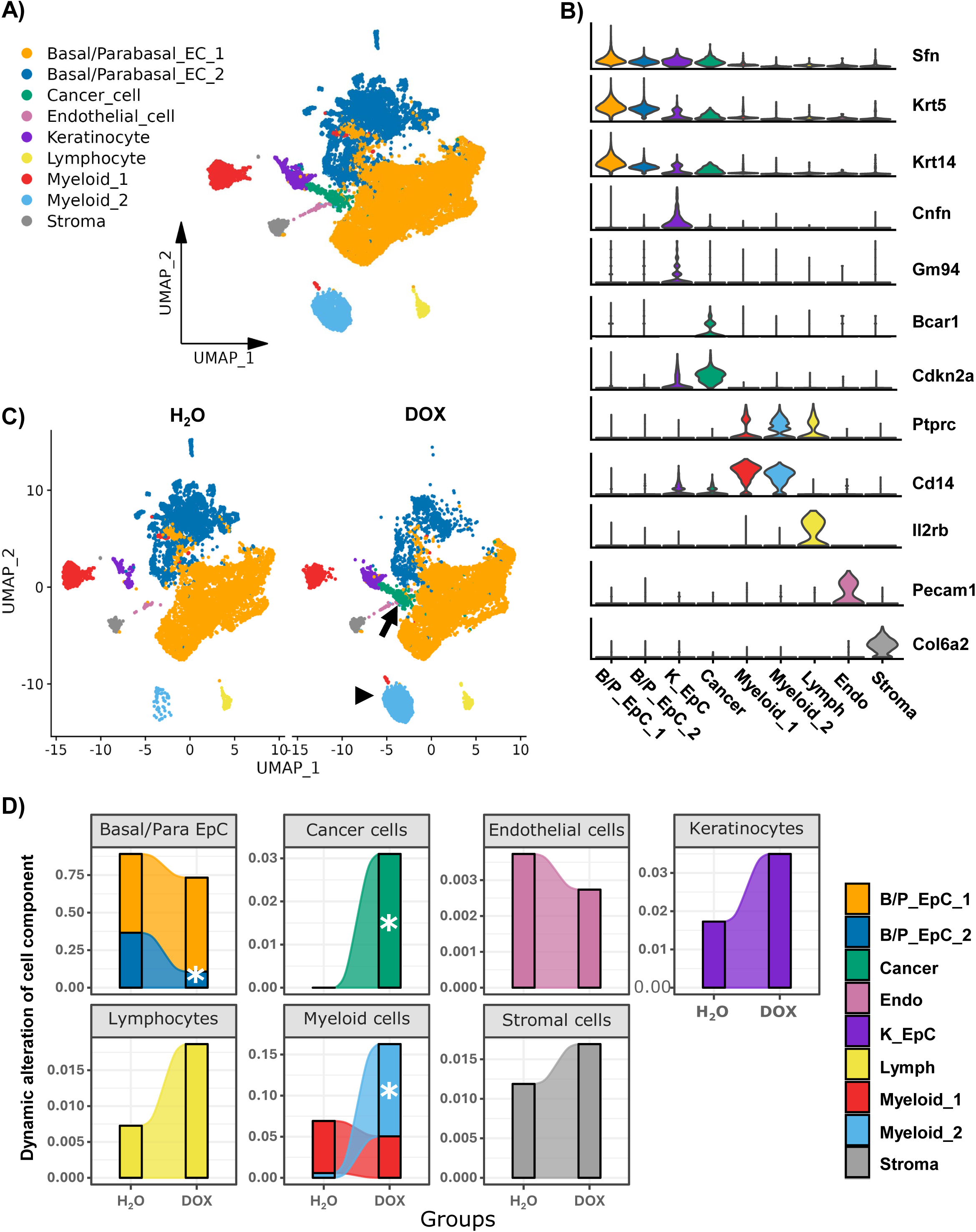
Single-cell RNA-seq reveals dynamic remodeling of the cervical cellular landscape during early tumorigenesis. **A**) UMAP plot showing overview of 9 major cell types identified in cervical tissues in *Krt14*-rtTA;Tet-on-YAP^S127A^ mice administered with water (H2O) as control or doxycycline (DOX) for tumorigenesis. **B**) Violin plots showing expression of marker genes for the identified cell clusters, including markers for Basal/Parabasal epithelial cell (EC) 1, Basal/Parabasal EC 2, Keratinocyte, Cancer cell, two myeloid cells (Myeloid_1 & Myeloid_2), Lymphocyte, Endothelial cell, and Stroma cells. **C**) UMAP plots showing the alteration of cellular components in cervical tissues of *Krt14*-rtTA;Tet-on-YAP^S127A^ mice administered with water (H2O) as control or doxycycline (DOX) for induction of tumorigenesis. Arrow points to the newly formed carcinoma cells, while the arrowhead points to the accumulated myeloid 2 cells (tumor-associated macrophages) during early-stage of tumorigenesis. **D**) Bar plots overlaid with a smoothed area curve showing dynamic alterations of major cellular components of cervical tissues of *Krt14*-rtTA;Tet-on-YAP^S127A^ mice during Doxycycline (DOX)-induced tumorigenesis. *Krt14*-rtTA;Tet-on-YAP^S127A^ mice treated with water (H_2_O) were used as control. *: significantly different from the control (H_2_O). Statistical differences were calculated with scCODA with FDR = 0.05.

All epithelial cell clusters, including the tumor cluster, expressed canonical epithelial markers *Sfn*, *Krt5,* and *Krt14*, consistent with their epithelial origin (25). As expected, *Krt5* and *Krt14*, which mark cervical basal and parabasal epithelial cells (26), were most highly expressed in the basal/parabasal epithelial cell cluster (B/P_EpC) (Fig. 3B). In contrast, *Cnfn* and *Gm94*, established markers of well-differentiated, keratinized superficial epithelial cells in the cervix (26), specifically labeled the keratinocyte cluster (K_EpC). *Bcar1* (p130Cas), a scaffolding protein that cooperates with mutant TP53 to promote invasion and motility (27), and *Cdkn2a*, which encodes the cervical cancer marker p16^INK4A (28), uniquely marked the tumor cluster (Fig. 3B). Three immune cell groups were distinguished by the pan-leukocyte marker *Ptprc* (CD45), including two myeloid populations characterized by *Cd14*, a marker of monocyte/macrophage lineage cells (29), and a lymphocyte cluster expressing *Il2rb*, a marker of T-cell activation and proliferation (30) (Fig. 3B). Endothelial and stromal clusters were confirmed by selective expression of *Pecam1* (CD31) and *Col6a2*, respectively (Fig. 3B).

Cancer cells were absent in cervical tissues from KY mice receiving normal water but emerged prominently in mice treated with doxycycline for four weeks, constituting ∼3% of all captured cells (FDR < 0.05; Fig. 3C-3D). This observation is consistent with the histopathological and immunohistochemical evidence of invasive carcinoma shown in Fig. 2. Among immune populations, one myeloid subset (Myeloid_2) also expanded significantly in doxycycline-treated mice (FDR < 0.05), suggesting recruitment and/or activation of this population during early tumorigenesis. A subset of basal/parabasal epithelial cells (B/P_EpC_2) decreased significantly upon YAP1 activation, indicating that this population may represent a cellular intermediate, or potential cell of origin, in the progression toward cervical cancer (Fig. 3C-3D).

### Spatial transcriptomics reveals spatially resolved cellular and molecular changes during YAP1-driven cervical carcinogenesis

To characterize in situ gene expression changes during cervical cancer development, we performed high-resolution spatial transcriptomics (Curio Seeker platform; 0.3 × 0.3 cm capture area; 10 µm spatial resolution, Curio Biosciences) on cervical tissue sections from KY mice treated with Dox or water (control). The analyzed sections matched those used for scRNA-seq, enabling integrative single-cell and spatial analysis. Spatial transcriptomic data preserved tissue architecture with high fidelity, recapitulating the histological features of adjacent H&E-stained sections (right panels, Fig. 4A-4B). The color-coded unsupervised clustering of spatial transcriptomic profiles identified major epithelial compartments, including basal/parabasal and superficial epithelial layers, in both samples (Fig. 4A - 4B). In the Dox-treated sample, the high-dimensional transcriptomic profile also delineated the tumor region (Fig. 4B), consistent with histopathological and IHC findings (Fig. 2).

**Figure 4.**
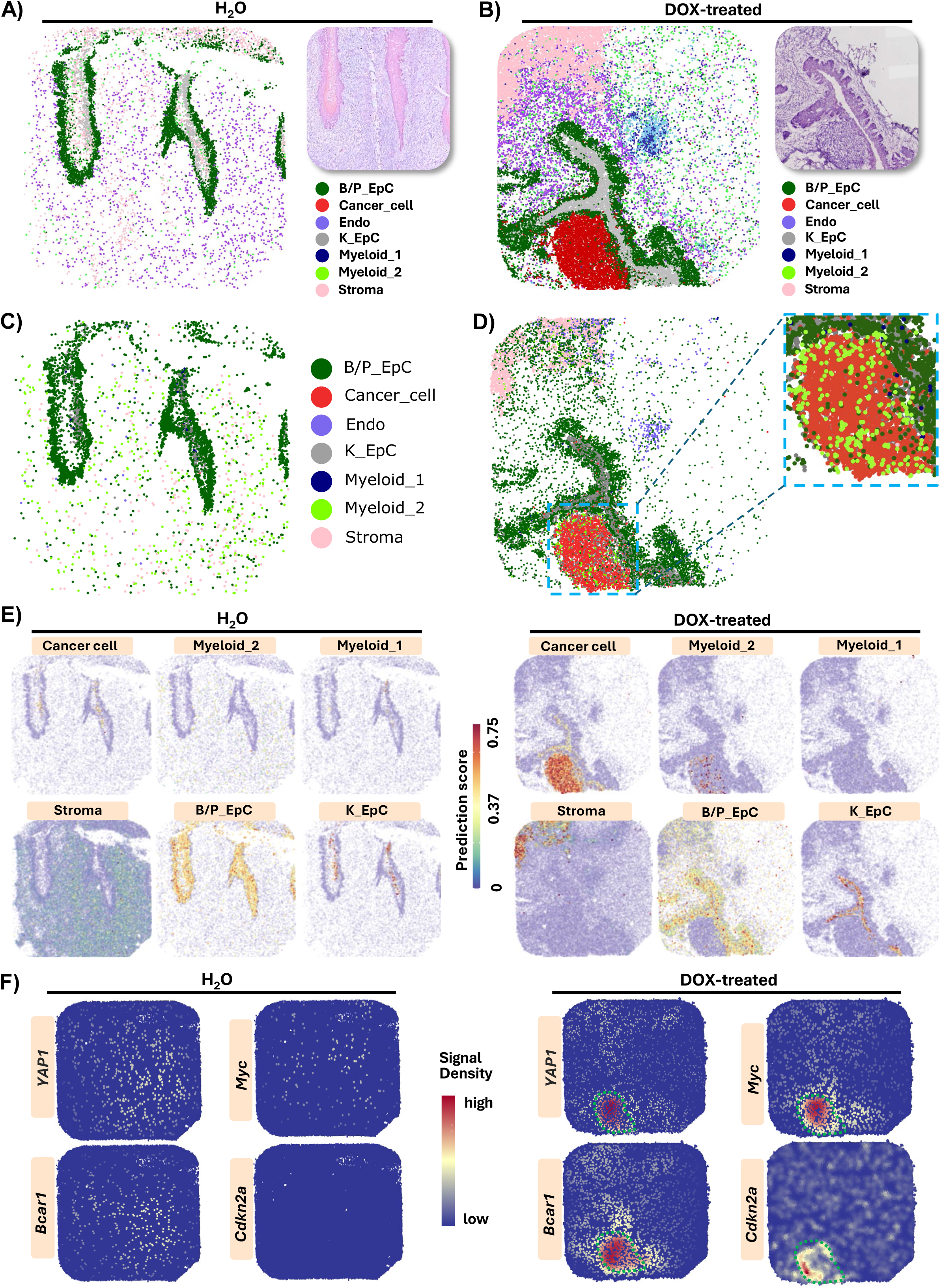
Mapping cervix cellular and molecular reprogramming during mesenchymal transition and invasion using spatial transcriptomics. **A & B**) Spatial transcriptomic plot showing that unsupervised clustering based on spatial transcriptomics data successfully captured the morphology of cervical tissue in *Krt14*-rtTA;Tet-on-YAP^S127A^ mice administered with water (H_2_O, panel A) and Doxycycline (DoX, panel B). The identified cell clusters are presented in different colors. **C & D**) Spatial distribution of sc-RNAseq-identified cells in the cervical tissues of KY mice treated with water (H_2_O, panel C, as negative control) or doxycycline (DOX, panel D). The cells were identified using marker/reference genes derived from the SC-RNAseq analyses of the matched samples. The color codes used to annotate cells in panel C) and panel D) are same. The dash-border square highlights the area with the newly formed cancer cells. A zoom-in view was presented on the right to visualize cancer cells (red) and tumor-associated myeloid cells (bright green, myeloid_2) in the tumor region. **E**) Spatial transcriptomic plots mapping the location of each individual cell type in cervical tissues in *Krt14*-rtTA;Tet-on-YAP^S127A^ mice treated with water as control (H_2_O, left) or doxycycline (DOX, right) to induce tumorigenesis. Cells were annotated using cell markers/references identified using sc-RNAseq analyses of the matched samples. Color intensity encodes the prediction score, corresponding to the confidence level of the cell-type assignment. **F**) Spatial transcriptomic plots mapping the expression and location of *YAP1*, *Myc*, *Bcar1* and *Cdkn2a* genes in cervical tissues of *Krt14*-rtTA;Tet-on-YAP^S127A^ mice treated with water as control (H_2_O, left) or doxycycline (DOX, right) to induce tumorigenesis. The dash-border circle highlights the cancer region. Sequential color scales represent relative expression levels of dedicated genes.

To leverage the complementary strengths of scRNA-seq (cellular precision, high transcript capture) and spatial transcriptomics (spatial resolution), we transferred cell-type annotations from the scRNA-seq dataset (Fig. 3) to the spatial transcriptomic spots using established integration workflows (24). Because both datasets were generated from matched tissues, cell-type transfer produced high-confidence spatial maps of the same biological cell types across modalities. The resulting cell-type-resolved spatial landscapes (Fig. 4C-4D) revealed that scRNA-seq-identified cancer cells (“Cancer_cell,” red) localized beneath the basal/parabasal epithelial layer (“B/P_EpC,” dark green) in the doxycycline-treated sample (Fig. 4D), consistent with the invasive growth pattern observed above.

To further visualize these cell-type assignments, we examined cell-type prediction scores, where higher scores reflect higher confidence (Fig. 4E–F). In the Dox-treated tissue, tumor regions exhibited strong and spatially coherent prediction scores for cancer cells. Notably, these invasive regions were infiltrated by the scRNA-seq–defined Myeloid_2 population, an observation evident both in the cell-type map (Fig. 4D) and in the layer-specific prediction scores (Fig. 4F). This finding corroborates the significant expansion of the Myeloid_2 cluster observed in scRNA-seq (Fig. 3C-3D) and suggests that this subset of myeloid cells may contribute to tumor initiation or progression.

Spatial maps also confirmed the presence of distinct epithelial layers, basal/parabasal epithelial cells and keratinocytes (“B/P_EpC” and “K_EpC”), in both samples (Fig. 4E). These findings support our interpretation that the KY model reflects an HPV-independent CVC subtype that develops beneath an intact epithelial surface and therefore may evade early detection by Pap smear screening. Finally, spatial transcriptomics validated the elevated expression of CVC-associated genes in the Dox-treated sample compared with controls, including *YAP1* and *Myc* (Fig. 4F), which were previously detected by IHC (Fig. 2), as well as *Bcar1* and *Cdkn2a* (Fig. 4F), identified in the tumor cluster of the scRNA-seq dataset (Fig. 3).

### scRNA-seq combined with spatial transcriptomics unveils the molecular mechanisms underlying the invasiveness of YAP1-induced cervical carcinoma cells

We then integrated scRNA-seq and spatial transcriptomics to uncover the molecular mechanisms underlying epithelial cell transformation and cancer cell invasiveness. In the scRNA-seq dataset, the cancer cell cluster demonstrated high-density *YAP1* expression among all identified cell clusters (Fig. 5A), consistent with the spatial transcriptomics data in which *YAP1*-expressing cells localized to the tumor region (Fig. 4E&4F). Quantitatively, the YAP1 SIGNATURE score was significantly enriched in cancer cells from DOX-treated samples relative to epithelial cells from H_2_O control group (Fig. 5B, supplementary Fig. S1). The significant decrease of B/P_EpC2 cluster and the development of cancer cells in cervical tissues of Dox-treated KY mice suggested that a portion of cancer cells may be derived from transformation of B/P_EpC2 cluster (Fig. 5C). Gene set enrichment analysis (GSEA) (31) was used to identify genes and pathways involved in the cell transformation during early stage of YAP1-induced cervical carcinogenesis. We found that the predominantly enriched genes and pathways during YAP1-induced tumorigenesis are those involved in epithelial to mesenchymal transition (EMT) and EMT-associated multicancer invasiveness, followed by macrophage migration and M2 macrophage-related negative regulation of immunity (Fig. 5D, Supplementary. Fig. S2). To validate EMT enrichment in situ, we performed Gene Set Variation Analysis (GSVA) (32) on the spatial transcriptomic dataset and visualized the density of GSVA-weighted EMT scores. EMT signature activity was strongly localized to the tumor region of the cervical tissues from the DOX-induced mice (Fig. 5E) but absent from adjacent non-tumor areas or any control (H_2_O-treated) tissues. These data suggest that EMT of transformed cervical epithelial cells and immunosuppressive microenvironment created by tumor-associated myeloid cells (TAMCs) may provide a foundation for the development of YAP1-induced invasive cervical cancer in Dox-induced KY mice. The clinical relevance of our finding is verified by the observation that 95% of signature genes of the EMT cluster identified in TCGA studies (17) are upregulated in cancer cells when compared to cervical epithelial cells in H_2_O-treated control groups (Fig. 5F, supplementary. Fig. S3). Consistently, around 86% of signature genes of human EMT cluster are upregulated when compared with non-malignant cervical epithelial cells in Dox-treated KY samples (supplementary. Fig. S4).

**Figure 5.**
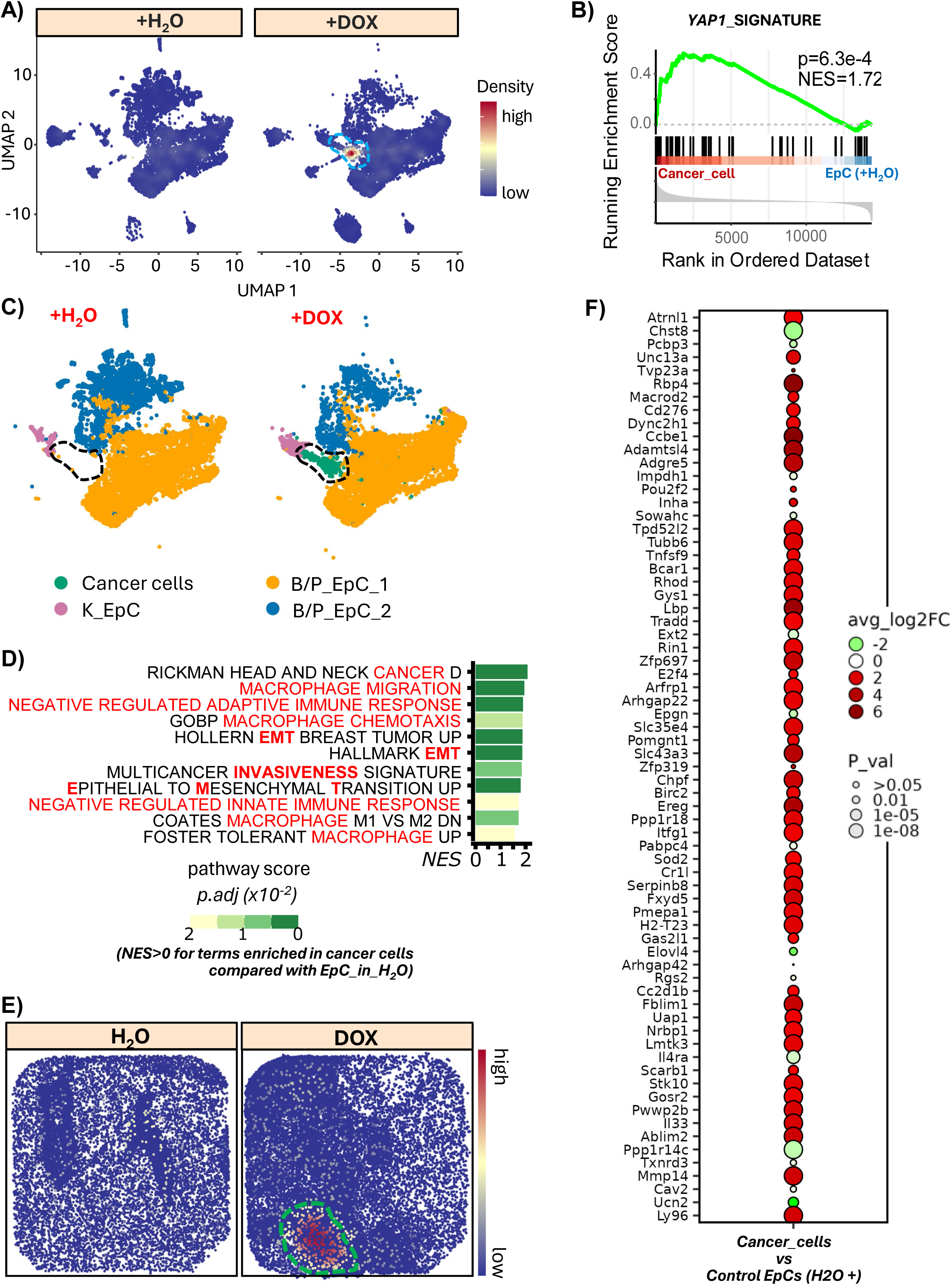
A combination of Single-cell RNAseq with spatial transcriptomics uncover the molecular mechanisms underlying the invasiveness of YAP1-induced carcinoma cells. **A**) UMAP plot derived from scRNAseq data showing density of YAP1 expression in cervical cells of KY mice administered with water as control (H_2_O) or Dox to induce cervical cancer. The dash-border circle highlights the cluster of cancer cells, which have the highest *YAP1* transgene. **B**) GSEA plot derived from scRNAseq data showing enrichment of YAP1 signature in cancer cells in *Krt14*-rtTA;Tet-on-YAP^S127A^ mice induced with doxycycline (DOX) compared with cervical epithelial cells in control mice (KY mice treated with H_2_O). **C**) UMAP plot derived from scRNAseq data showing epithelial cell subtypes in cervical tissues in *KY* mice treated with water (H_2_O) as control or Dox to induce CVC. The dash-border circle highlights the cluster of cancer cells. B/P_EpC; basal/parabasal epithelial cells; K_EpCs: Keratinized cervical epithelial cells. **D**) The pathways enriched in invasive cancer cells in Dox-induced KY mice when compared to cervical epithelial cells in control mice (KY mice treated with H2O as negative control). Color scales show adjusted *P* values. **E**) Spatial transcriptomics plot showing density of EMT signature (77) subtypes in cervical tissues in control (H_2_O) and Dox-induced KY mice. The dash-border circle highlights the cluster of cancer cells. Sequential color scale represents the signal density of EMT signature genes. The dash-border circle highlights the cancer region. **F**) A bubble chart showing the upregulation of EMT signature genes in invasive cancer cells. The relative gene expression data of EMT signature genes in invasive cancer cells and control cervical epithelial cells were derived from single-cell RNA sequencing analyses of single cells isolated from cervical tissues from doxycycline-induced KY mice and non-induced KY mice (H_2_O+, used as negative control). The color represented average log2 fold changes (avg_log2FC) in gene expression levels of invasive cancer cells relative to non-induced control cervical epithelial cells (control EpCs-H2O+). The size marks the p values generated by non-parametric Wilcoxon rank sum test implemented in R package Seurat. The analysis was restricted to human EMT signature ortholog genes (17). Notably, 95% of EMT signature genes are upregulated in invasive cancer cells compared to non-induced control cervical epithelial cells.

### scRNA-seq combined with spatial transcriptomics reveals mechanisms by which YAP-high cancer cells create an immunosuppressive microenvironment

Accumulation of Myeloid cells is another major feature of YAP1-induced invasive CVC (Fig. 4D–F). To characterize the myeloid population intermixed with cancer cells, we performed higher-resolution clustering of myeloid cells (24). Unsupervised analysis, supported by established markers, resolved three distinct subsets: myeloid-derived suppressor cells (MDSCs), tissue-resident macrophages (TRMs), and a third cluster containing fewer macrophages (mono/M) (Fig. 6A-6B). All three subsets expressed canonical myeloid markers *Lyz2* and *Cd14* (Fig. 6A-6B). Low expression of *Cd68* and *Mpeg1*, two pan-macrophage markers, distinguished the mono/M population (Fig. 6A, 6B). TRMs were identified by high expression of immunosuppressive markers *Apoe* and *Spp1* (33) and type II macrophage markers *Ddit4* and *Il4ra* (34, 35) and by their stable abundance irrespective of tumorigenesis (Fig. 6A–B, supplementary Fig. S5). In contrast, the MDSC cluster was defined by expression of immunosuppressive mediator *Il10ra* together with MDSC-associated markers *Cd84* and *Bcl2l1* (36, 37). Low *Ddit4* expression further indicated that these MDSCs were distinct from type II macrophages. Importantly, we found that the observed myeloid cell accumulation could be attributed to the significant increase in the number of MDSCs (Fig. 6C).

**Figure 6.**
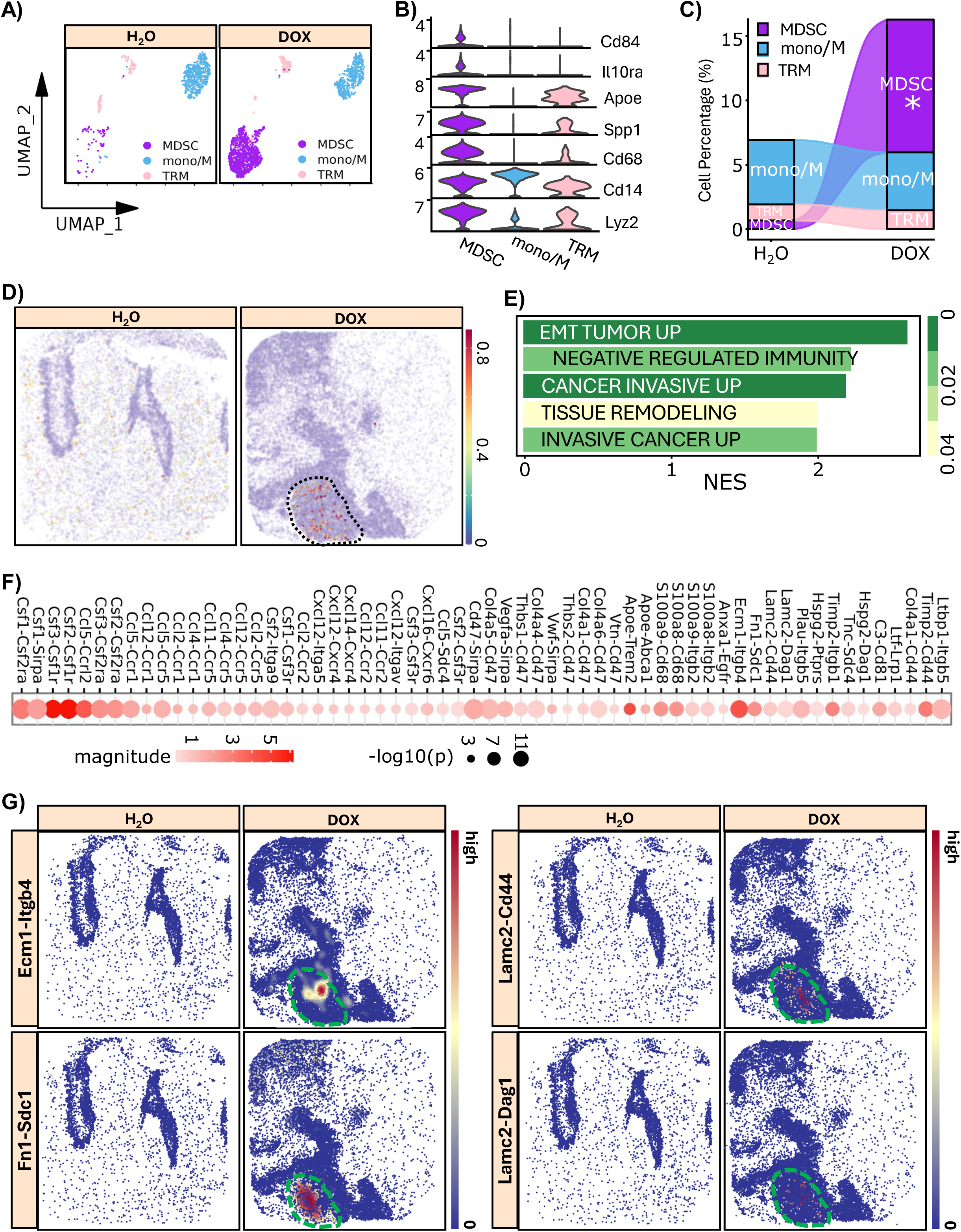
A combination of Single-cell RNAseq with spatial transcriptomics reveals a role of MDSC in mesenchymal cervical cancer development. **A**) UMAP plot showing subtypes of myeloid cells in cervical tissue in *Krt14*-rtTA;Tet-on-YAP^S127A^ mice administered with water as control (H_2_O) or doxycycline (Dox) to induce cervical cancer. MDSC: Myeloid-derived suppressor cells; Mono/M: other myeloid cells; TRM: Tissue resident macrophages. **B**) Violin plots showing expression of representative marker genes of the identified the major myeloid subtypes in A). **C**) Dynamic alterations of myeloid cell subtypes in cervical tissues in control (H_2_O) and doxycycline-induced (DOX) Krt14-rtTA;Tet-on-YAPS^127A^ mice. *: significantly different when compared to control group. Statistical difference was performed with scCODA with FDR= 0.05. **D**) Spatial transcriptomic plot showing the location of MDSCs (identified using matched scRNAseq samples as reference) subtypes in cervical tissues in control (H_2_O) and doxycycline-induced (DOX) *Krt14*-rtTA;Tet-on-YAPS^127A^ mice. Sequential color scales represent the prediction score / confidence of cell type assignments. The dash-border circle highlights the cancer region. **E**) The pathways enriched in MDSCs in the cervical tissues of *Krt14*-rtTA;Tet-on-YAPS^127A^ mice induced with doxycycline (DOX) when compared with that of myeloid cells in cervical tissues of the control mice (*Krt14*-rtTA;Tet-on-YAPS^127A^ mice administered with H_2_O). Color scales show adjusted P values. **F)** The active ligand-receptor interactions between cancer cells and MDSCs. Magnitude is calculated as the arithmetic average of the mean expression of the ligand in the senders and the receptor in the receivers, using the minimum subunit expression for any multi-subunit complexes involved. *P* values were derived by random permutation of cell cluster labels implemented in Python package CellphoneDB. A *p*<1E-3 is considered specific. Please note that the primary interactions are associated with chemotaxis, immunosuppression, EMT and tumor invasiveness.**G)** Spatial distance and signal strength weighted density of ligand-receptor interactions between cancer cells and MDSCs in cervical tissues of control (H_2_O) and doxycycline-induced (DOX) *Krt14*-rtTA;Tet-on-YAPS^127A^ mice. Please note that the primary interactions are associated with immunosuppressive, EMT and tumor invasive. Sequential color scales represent the signal density of a specific interaction between cancer cells and MDSCs in the indicated tissues.

Next, we transferred these annotations to matched spatial transcriptomic datasets (24). MDSCs were markedly enriched within the tumor region (Fig. 6D), whereas TRMs and mono/M cells remained sparse in cervical tissue across both Dox-treated and H_2_O control groups (supplementary. Fig. S5), consistent with scRNA-seq findings.

We then performed GSEA to assess the functional contribution of MDSCs in tumorigenesis, and found that EMT-tumor, immunosuppression, and cancer invasiveness gene sets were significantly enriched in MDSCs from Dox-treated group compared with myeloid cells from H_2_O-treated control group (Fig. 6E). Cell-cell interaction analysis further uncovered multiple ligand-receptor pairs connecting cancer cells and MDSCs. These included immunosuppressive interactions (*C3–Cd81*, *Plau–Itgb5*, *Fn1–Sdc1*, *Anxa1–Egfr*, *S100a8/9*, *Apoe*, and *Cd47*) (33, 38–43), tumor-invasive interactions (*Ltbp1–Itgb5*, *Col4a1–Cd44*, *Ltf–Lrp1*, *Tnc–Sdc4*, *Lamc2–Dag1*, *Lamc2–Cd44*, and *Ecm1–Itgb4*) (40, 44–49), and chemotactic signals involving *Cxcl*, *Ccl*, and *Csf* family ligands (Fig. 6F).

Using our spatial transcriptomic datasets, we visualized the density of immunosuppressive and tumor-invasive ligand-receptor interactions, weighted by spatial distance and signal strength, specifically between cancer cells and MDSC (Fig. 6G). Notably, several identified interactions have well-established tumor-promoting functions. *Ecm1–Itgb4* drives metastasis via integrin-FAK/SOX2/HIF-1α signaling (40). *Sdc1*, a surface marker of intratumoral MDSCs, enhances tumor invasion via *Fn1–Sdc1*–mediated ECM remodeling and immune modulation (41). *Lamc2* promotes metastatic progression in lung adenocarcinoma (48), and *Lamc2–Cd44* interactions stimulate migration of metastatic breast cancer cells (49).

## DISCUSSION

November 17, 2025 marks the fifth anniversary of the WHO CVC elimination initiative. While the global progress enabled by Pap testing, HPV-based screening, and HPV vaccination is remarkable, these strategies rely on the long-standing assumption that all cervical cancers arise from HPV infection. Our previous studies, however, demonstrated that a subset of CVCs develops through an HPV-independent pathway (16). Together with the long-recognized limitations in Pap test sensitivity (50–53), these observations raise the possibility that current elimination efforts may overlook important biological and clinical gaps. Indeed, cervical cancer still causes substantial mortality worldwide (14, 54), and CVC deaths in the United States have remained essentially unchanged over the past two decades (15).

Although most cervical tumors harbor HPV DNA, multiple lines of evidence indicate that HPV alone is insufficient to induce malignancy. In humans, the overwhelming majority of HPV-positive epithelial cells are eliminated within 1–2 years of infection (55–58). In transgenic mouse models, expression of HPV E6 and/or E7 failed to induce cervical cancer in the absence of high-dose estrogen supplementation (59–61). While HPV is often considered “necessary,” emerging data call this into question. We previously showed that hyperactivation of YAP1 is sufficient to initiate cervical cancer in vivo, independent of HPV (16). With deep sequencing technologies, previous studies have also identified “fossil” HPV DNA, detectable viral genomes without transcriptional activity, in human cervical tumors, suggesting that DNA positivity does not guarantee causal involvement. Large datasets, including TCGA, report that ∼5–10% of cervical cancers lack molecular evidence of HPV-driven transcription (17, 62–65). Importantly, the true percentage of HPV-independent tumors may be underestimated, because epidemiologic studies do not account for post-neoplasia HPV infection. Our recent work shows that transformed cervical cancer cells are more susceptible to HPV infection than primary epithelial cells (66), consistent with earlier findings that proliferative cells are more readily infected (67). Similarly, although HPV detection increases with lesion grade (CIN1 < CIN2 < CIN3 < cancer), only ∼40% of CIN1 lesions are HPV-positive (63), indicating that a substantial proportion of early lesions develop without HPV involvement. Collectively, these data support the existence of bona fide HPV-independent cervical cancers and argue that HPV is not universally required for CVC initiation. Current screening approaches may be particularly ineffective for detecting HPV-independent tumors.

The Pap test, since its introduction in the mid-20th century (3, 68), has substantially reduced CVC rates in countries where this technology is widely implemented. However, large-scale studies demonstrate that its sensitivity may be <50% (50–53, 69). The low sensitivity was generally attributed to the low test performance such as inadequate sample collection, poor quality of sample, improper timing of test, and even the inexperience of health provider. Our results provide a mechanistic explanation: YAP1-hyperactivated tumor cells adopt an early mesenchymal/invasive phenotype, enabling them to rapidly breach the basement membrane and invade into the cervical stroma, escaping exfoliative sampling. Thus, Pap test insensitivity likely arises not only from technical limitations but also from an incomplete assumption that all cervical cancers grow toward the epithelial surface.

Our spatial transcriptomics and single-cell analyses further define a YAP1-high EMT-subtype of CVC characterized by disrupted Hippo signaling, early accumulation of myeloid-derived suppressor cells (MDSCs), and extensive extracellular matrix remodeling. These features reveal therapeutic vulnerabilities. In papillary renal cell carcinoma, inhibitors targeting the YAP1–TEAD interaction (e.g., MGH CP-1) effectively prevented tumor initiation (23, 70, 71). Similar strategies may suppress or prevent YAP1-associated EMT-subtype of cervical cancers. Moreover, early MDSC accumulation suggests that myeloid targeting and MDSC-based biomarkers may enable early detection of this subtype.

In summary, our findings reveal a biologically distinct, YAP1-high subtype of cervical cancer that is neither reliably detected nor prevented by pap tests- and HPV-based strategies. By integrating single-cell and spatial transcriptomics, we identified cellular and molecular, and microenvironmental features that can guide the development of Hippo signaling pathway-based biomarkers, diagnostic approaches, and preventive therapeutics. Recognizing and addressing HPV-independent and YAP1-driven invasive CVCs will be essential for achieving true global cervical cancer elimination.

## MATERIALS AND METHODS

### Analyses of Human cervical cancer patient genetic/genomic data

Genetic, genomic, and clinical data for all patients were sourced from The Cancer Genome Atlas (TCGA) CVC dataset (PanCancer Atlas) (17, 18). The dataset includes data from 297 cases, with 112 cases further stratified into epithelial–mesenchymal transition (EMT), hormone, or PI3K-Akt subtypes. Survival analysis for CVC was performed using the cBioPortal for Cancer Genomics (available at http://www.cbioportal.org) and visualized with R packages *survminer* (v0.4.9) and *survival* (v3.7.0) (72, 73). Violin, pie, and dot plots were generated using the R package *ggplot2* (v3.5.1). Copy number alterations (CNAs) significantly different across subtypes were identified using cBioPortal (http://www.cbioportal.org) (72, 73). YAP1 signature scores were calculated by summing the z-scores of genes associated with the YAP1 signature as described previously (17).

### Generating genetically modified mouse models

All mouse handling and experimental procedures were approved by the Institutional Animal Care and Use Committee (IACUC) at Massachusetts General Hospital (MGH) and University of Nebraska Medical Center (UNMC). Krt14-rtTA mice (FVB-Tg(KRT14-rtTA) F42Efu/J, #008099, The Jackson Laboratory) were crossed with Tet-on-YAP^S127A^ mice (a gift from Dr. Fernando Camargo’s lab, Boston Children’s Hospital) to generate Krt14-rtTA; Tet-on-YAP^S127A^ mice, which express a constitutively active form of YAP1 (YAP^S127A^) under the control of a Krt14-driven tetracycline regulatory element (TRE) (74). Genotyping was performed using RT-PCR on tail tissue with primers listed in Table S1. Transgenic mice older than 2 months were implanted subcutaneously with 17β-estradiol pellets (0.25 mg/pellet, 90-day release, Innovative Research of America Inc.) and treated with 0.03 mg/mL doxycycline (DOX) in drinking water to induce transgene expression.

### Immunohistochemistry

YAP1, Myc, Ecad and VIM in both normal and cancerous cervical tissues were detected using an established immunohistochemistry (IHC) protocol in our lab (66). The antibodies used for IHC are detailed in Table S2. Immunosignals were visualized using a polymer-based immunohistochemistry kit and a DAB assay kit (Vector Laboratories, Burlingame, CA). The stained slides were scanned and imaged using an MorphoLens 1 Scanner (Morphle Labs Inc., NY).

### Single-cell Transcriptomics and Bioinformatic Analyses

Cervical tissues from control and transgenic mice were collected and processed using a protocol described by Chumduri et al. (25). Briefly, cervical tissues were excised, washed in sterile PBS (Gibco, 14190-094), and minced into small pieces (∼1 mm²) on ice. The tissue was then digested at 37°C for 40 minutes in an air bath shaker set to 200 rpm. The digestion medium consisted of 450 units/mL collagenase I, 0.8 units/mL elastase, 300 units/mL hyaluronidase, 250 units/mL DNase I, and 1 unit/mL dispase. After digestion, the cell suspension was mixed with cold RPMI 1640 supplemented with 2% FBS, sequentially filtered through pre-wetted 70-μm and 40-μm strainers, and centrifuged at 1500 rpm for 5 minutes at 4°C. The resulting cell pellets were subjected to ACK lysis for 10 minutes to remove red blood cells. For single-cell sequencing, cells isolated from 5–6 mice within the same group were pooled, and each tissue was processed separately. The prepared cells were assessed for viability, counted, and adjusted to the appropriate concentration for single-cell RNA sequencing library preparation.

Single cells were partitioned into nanoliter-scale Gel-Bead-In-Emulsions (GEMs) using a 10x Chromium Controller. Single-cell libraries were then processed using the 10x Genomics Chromium Single Cell 5′ Library & Gel Bead Kit v2 (PN-1000014), targeting 10,000 cells per channel. Reverse transcription was performed using a C1000 Touch Thermal Cycler (Bio-Rad Laboratories). Library quality and quantification were assessed using the Agilent 2100 Bioanalyzer with a DNA High Sensitivity chip. Libraries were sequenced on an Illumina HiSeq 2500 NexGen Sequencer, with approximately 500 million reads allocated per sample, and around 50,000 reads per cell.

All FASTQ files passed standard sequencing quality control metrics and were processed using 10x Genomics Cell Ranger. The resulting raw counts for each sample were evaluated for quality control using Seurat (version 5.0). Each Seurat object was normalized using the SCTransform algorithm (version 2) and integrated within Seurat. Differential gene expression analysis was performed using Wilcoxon tests. Clustering was based on the Jaccard similarity of the K-nearest neighbors (KNN) graph, refined using Euclidean distance in PCA space. These analyses and visualizations were conducted using Seurat and SCpubr (version 1.0.4). For quality control, isolated cells were filtered to retain only those with < 20% mitochondrial gene expression, > 500 RNA counts, and > 170 RNA features. Potential doublets were identified and excluded using Scrublet (from the scran package, version 1.14.6).

### Spatial transcriptomics

Spatial transcriptomics was performed on cervical tissues that were matched with those used for single-cell RNA sequencing, employing the Curio Seeker 3 × 3 kit according to the manufacturer’s protocol (CB-000470-UG v2, Curio Biosciences). Blood and other contaminants were carefully removed, and fresh tissues were immediately embedded in OCT media (SAKURA, 25608-930) and frozen in a liquid-nitrogen-cooled isopentane (EMD Millipore, MX0760) bath. The tissue blocks were then sectioned into 10-µm slices using a Leica CM1950 cryostat and mounted onto Curio Seeker Tiles. A barcode whitelist and barcode position file corresponding to each tile were provided by Curio Biosciences. After RNA hybridization and reverse transcription, the tissue sections were digested, and the beads were removed from the glass slides and resuspended. Second-strand synthesis was performed using semi-random priming, followed by cDNA amplification. For the tagmentation, 1,200 pM DNA was used with the Nextera XT Library Prep Kit (FC-131-1024, Illumina). Libraries were pooled and sequenced on an Illumina HiSeq 2500 NexGen Sequencer. The sequenced data were processed using the Curio Seeker pipeline v.2.0.0 ([Curio Bioscience Bioinformatics Pipeline](https://curiobioscience.com/bioinformatics-pipeline)).

### Statistics and Reproducibility

All experiments were performed independently at least three times unless otherwise stated. Sample sizes, including those for animal models, were determined based on our previous studies (16, 23, 66, 75, 76). Samples were randomly assigned to experimental groups. Data are expressed as *mean* ± *SEM*. Statistical analyses were conducted using GraphPad Prism software (GraphPad Software, Inc., La Jolla, CA), or as otherwise specified. Significance was assessed using Student’s t-tests, one-way ANOVA, log rank test, Pearson’s Chi-squared test, non-parametric Wilcoxon rank sum test, or the tests implemented in R or Python packages such as fgsea, scCODA, CellPhoneDb as detailed when first mentioned. Differences were considered statistically significant when *P* < 0.05.

## ETHICS APPROVAL AND CONSENT TO PARTICIPATE

Transgenic mouse models were generated in this study to examine the role of hyperactivated YAP1 oncoprotein in the development of invasive cervical cancer in the transformation zone of uterine cervix. All animal model-associated experimental procedures were approved by the Institutional Animal Care and Use Committee (IACUC) of the University of Nebraska Medical Center (UNMC) and Massachusetts General Hospital (MGH). No human subjects were involved in this study.

## DATA AVAILABILITY

The raw and processed RNAseq data generated in this study have been deposited in the NCBI Gene Expression Omnibus (GEO) database under the accession number/code GSE315003 [https://www.ncbi.nlm.nih.gov/geo/query/acc.cgi?acc=GSE315003] and GSE315004 [https://www.ncbi.nlm.nih.gov/geo/query/acc.cgi?acc=GSE315004]. Other data supporting the findings of this study are available within the article and its supplementary information files.

## ACKNOWLEDGEMENT

This work was partially supported by the National Cancer Institute / National Institute of Health (1R01CA197976, 1R01CA201500, and 5R01CA279385), the Colleen’s Dream Foundation (no number), the Ruggles Family Foundation, and the Vincent Memorial Hospital Foundation / Vincent Department of Obstetrics and Gynecology, Massachusetts General Hospital (No number).

## AUTHOR CONTRIBUTIONS

J.L and X.L contributed to the conceptualization, experimental design and performance, data analysis, and manuscript preparation; X.L, J.L, C.H (Chunbo He), and G.H contributed to transgenic mouse model generation; X.L, C.H (cong Huang), R.J, and P.C contributed to transgenic mouse strain maintenance; J.L contributed to single cell isolation; R.S contributed to RNA sequencing; J.L contributed to visualizing large-scale data derived from single cell RNA-sequencing and spatially resolved transcriptomics; J.L contributed to Fluorescent immunohistochemistry (FL); X.L., J.L., and P.C. contributed to Immunohistochemistry (IHC); C.H (Cong Huang), P.C, X.Z, D.S, S.Y, M.M, B.C, A.P, G.K, R.G, A.M, and O.A conducted to experimental reagent preparation, real-time PCR (e.g., transgenic mouse genotyping), and histological analyses (e.g., tissues section, staining, and imaging, *etc.*); All authors contributed to manuscript review; J.D. contributed to result discussion and data interpretation; C.W. supervised these studies and contributed to conceptualization, experimental design, data analysis/interpretation, and manuscript preparation.

