## supplemental information for "Hyperactivated YAP1 Drives an Invasive EMT Subtype of Cervical Squamous Cell Carcinoma"

Cancer cells vs other cervical EpCs in Dox-induced KY mice

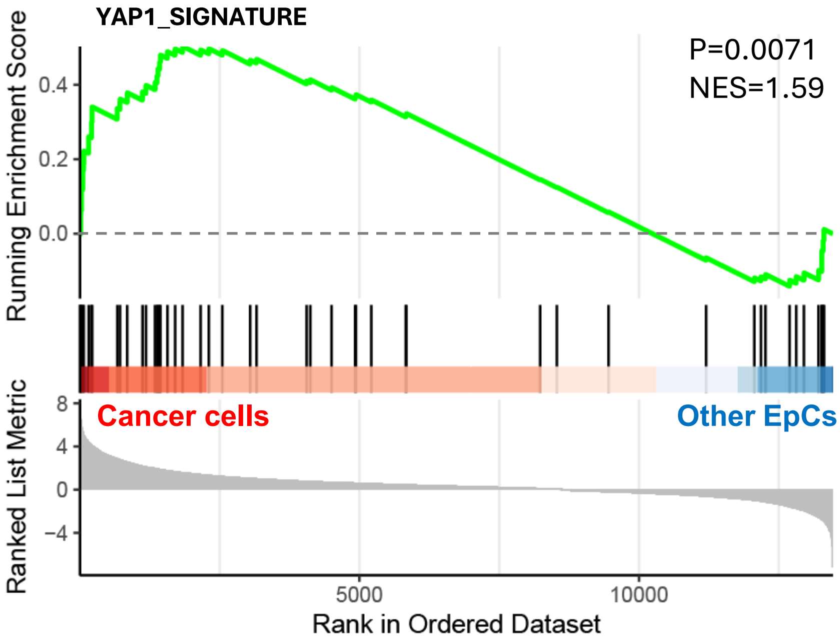


**Figure S1. GSEA plot showing enrichment of YAP1 signature genes in cancer cells and non-transformed epithelial cells (other EpCs) in cervical tissues of *Krt14*-rtTA;Tet-on-YAP$^{S127A}$ mice induced with doxycycline (DOX).** Gene expression data were derived from Single-Cell RNA sequencing. (*Relevant to Fig. 5B*)

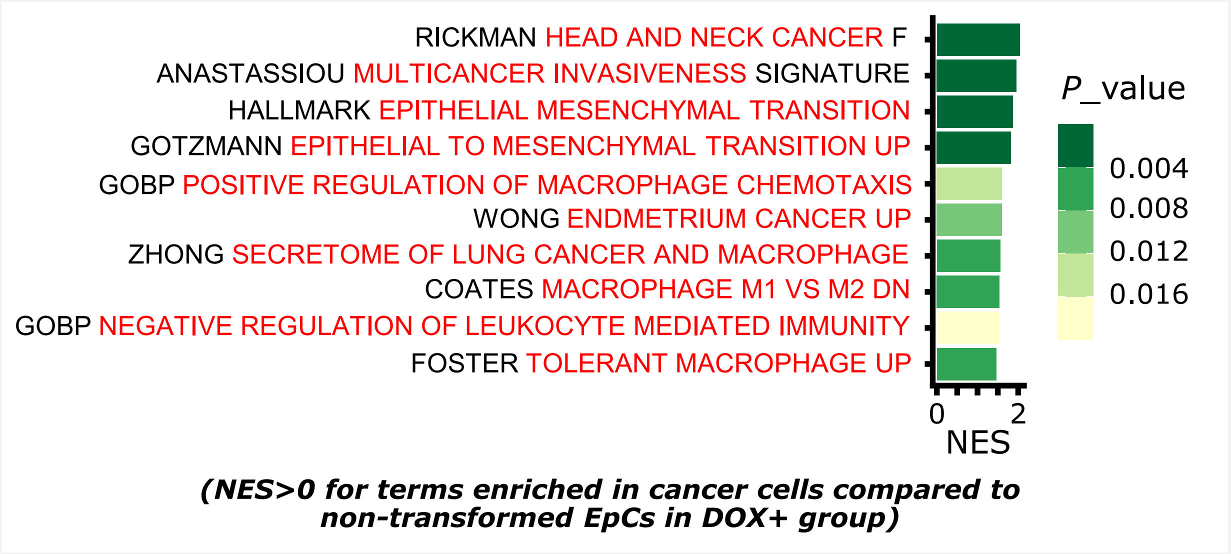


**Figure S2. Gene set enrichment analysis (GSEA) highlighting predominantly enriched pathways in YAP1-induced invasive cancer cells.** GSEA was performed using single-cell RNA sequencing data. The enrichment of genes in the invasive cancer cells was calculated by comparing them to genes in non-transformed cervical epithelial cells (Non-transformed EpCs) from the same Doxycycline-induced KY mice (Dox+ group). Notably, the predominantly enriched pathways during YAP1-induced cervical tumorigenesis include epithelial-to-mesenchymal transition (EMT) and EMT-associated multicancer invasiveness, followed by macrophage migration and M2 macrophage-related negative regulation of immunity. (*Relevant to Fig. 5D*)

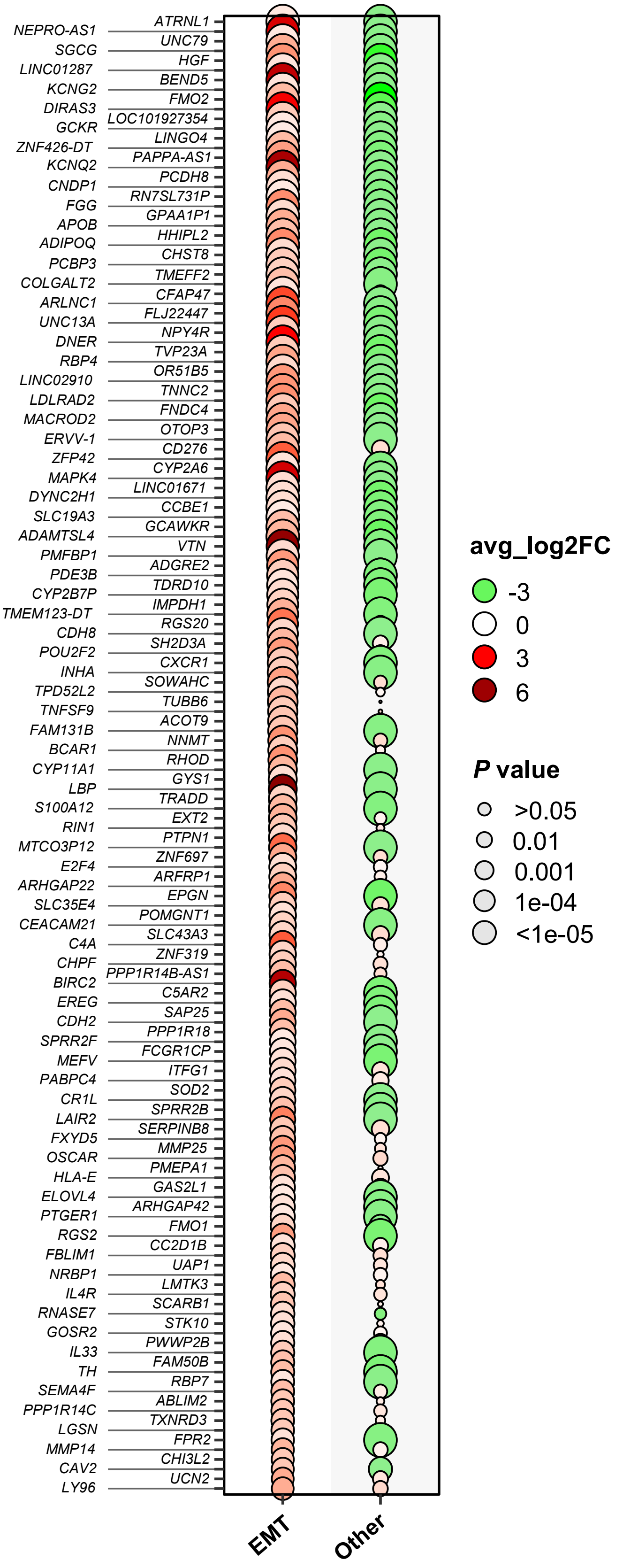


**Figure S3. A bubble chart illustrating the relative expression of cervical cancer subtype-specific genes in tumors derived from Dox-induced KY mice**. Gene expression data for mouse cervical cancer and normal cervical epithelial cells were obtained via single-cell RNA sequencing of cervical tissues from Dox-induced KY mice and normal control mice, respectively. The relative expression of signature genes for human epithelial-to-mesenchymal transition (EMT) and other subtypes (a combination of PI3K-AKT and Hormone subtypes) in mouse cancer cells is presented as the average $\log_2$ fold change (avg_$\log_2$FC) relative to that of the normal control. The analysis focused on human cervical cancer (CvC) subtype-specific ortholog genes (subtype-specific signature genes) expressed in mouse tumor cells. Notably, 95% of EMT subtype signature genes are upregulated in tumor cells derived from Dox-induced KY mice, while the signature genes for other CvC subtypes are not significantly upregulated. EMT: EMT cancer Vs Normal control; Other: other cancer Vs Control. (*Relevant to Fig. 5F*)

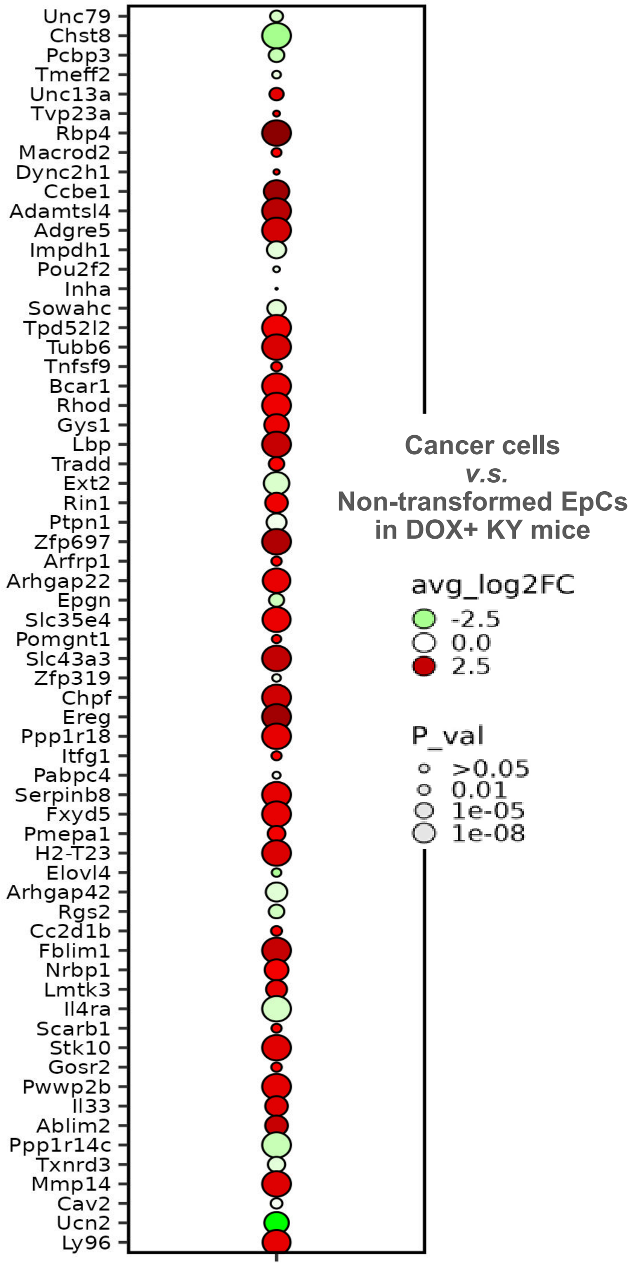


**Figure S4. A bubble chart illustrating the upregulation of EMT signature genes in invasive cancer cells.** The relative expression levels of EMT signature genes in invasive cancer cells and non-transformed cervical epithelial cells (non-transformed EpCs) from doxycycline-induced (Dox+) KY mice were derived from single-cell RNA sequencing analysis of cervical tissues with Dox-induced early CVC lesions. The data are presented as average $\log_2$ fold changes (avg_$\log_2$FC) in gene expression levels of invasive cancer cells relative to control non-transformed cervical epithelial cells (non-transformed EpCs) within the same tissue. The analysis focused on human EMT signature ortholog genes that were differentially expressed in mouse tumor cells. Notably, 86% of EMT signature genes are upregulated in invasive cancer cells compared to non-transformed cervical epithelial cells. (*Relevant to Fig. 5F*)

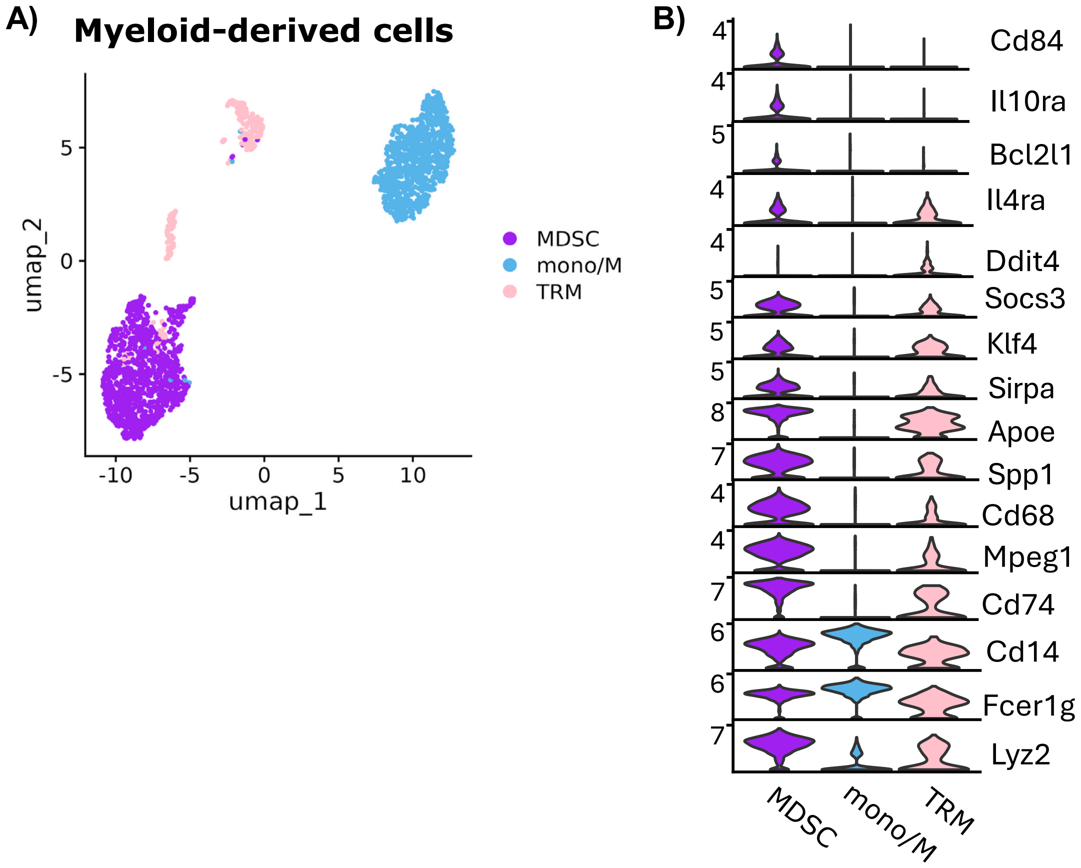


**Figure S5. Multiple subtypes of myeloid-derived cells are present in healthy and cancerous mouse cervical tissues. A)** A UMAP plot showing three major myeloid cell sub-clusters identified within myeloid-derived cells in healthy and cancerous mouse cervical tissues: MDSC (myeloid-derived suppressor cells), Mono/M (normal monocytes), and TRM (tissue-resident macrophages). **B)** Violin plots illustrating the relative expression levels of the whole panel of representative marker genes across the identified myeloid cell sub-clusters, as presented in Figures 6A–6C.
